# Characterization of heat shock protein expression and its application to temporal gene expression in the nematode *Pristionchus pacificus*

**DOI:** 10.1101/2025.06.24.661439

**Authors:** Yuuki Ishita, Hirokuni Hiraga, Takahiro Chihara, Misako Okumura

## Abstract

Temporal gene expression systems are widely used to examine gene functions at specific developmental stages. One method applied in temporal gene expression systems in many organisms is the heat-inducible gene expression system, which utilizes a heat shock promoter with evolutionarily conserved heat shock elements. The nematode *Pristionchus pacificus* is a satellite model system comparable to *Caenorhabditis elegans,* with unique developmental traits but lacking genetic tools for temporal gene expression. To establish a temporal gene expression system in *P. pacificus*, we investigated genes that were highly induced by heat shock. RNA-seq analysis revealed thousands of differentially expressed genes after a 2-hour heat shock event. One of the highly induced genes, *PPA12242*, is an ortholog of *C. elegans hsp-16.41*, and reporter transgenic animals have indicated that the genomic fragment upstream of this gene can induce gene expression in response to heat shock. *PPA12242* can be highly induced in specific larval stages, which seem to be vulnerable to heat stress in some phenotypes. Taken together, we identified a potential heat shock promoter in *P. pacificus* that is applicable to the temporal gene expression system in this species.

## Introduction

Temporal gene expression systems are powerful tools for investigating gene function in specific developmental stages in genetic model organisms. Over the past several decades, various methods for temporal gene expression and suppression have been developed, including RNA interference, recombinase-based expression systems, the Tet-On/OFF system, and the Gal4-UAS system (see (Driesschaert, Mergan, & Temmerman, 2021) for review). Using these techniques, the developmental roles of numerous genes have been examined not only in model systems but also non-model organisms (Abete-Luzi et al., 2020; Hari et al., 2012; Lee & Luo, 1999; Pavlopoulos et al., 2009; Watanabe et al., 2007). The heat-inducible gene expression system is one of the most widespread temporal gene expression methods. This system utilizes a heat shock promoter containing repeats of evolutionarily conserved heat shock elements with NGAAN and NTTCN motifs (Amin et al., 1988). Upon heat stress, the HSF1 transcription factor binds to the tandem repeats of heat shock elements, and stress-induced genes, such as those encoding molecular chaperones, are transcribed. Since the first report of the *Drosophila hsp70* gene (Pelham, 1982), heat shock promoters have been identified and utilized for heat-inducible gene expression in model organisms and some non-model organisms (Kawaguchi et al., 2015; Schinko et al., 2012; Xing et al., 2017).

The nematode *Pristionchus pacificus* is a satellite model nematode comparable to *Caenorhabditis elegans*, a classical genetic model animal (Sommer et al., 1996). With an apparent association with insects in the wild, numerous strains of this species, as well as over 40 new species of the same genus, have been discovered in the past few decades (Kanzaki et al., 2021). Since this species was first described in 1996 (Sommer et al., 1996), various genetic tools, including forward and reverse genetics, transgenic reporter lines, genetic cell ablation, and inhibition of neuronal transmission, have been developed (Han et al., 2020; Nakayama et al., 2020, 2024; Okumura et al., 2017; Rödelsperger et al., 2017; Schlager et al., 2009; Witte et al., 2015). Using these tools, the genetic mechanisms of various interesting features in this nematode have been characterized, such as the convergent evolution of vulval morphology, developmental plasticity of the mouth, evolution of predatory feeding behavior and associated mouth structure, kin recognition, and chemosensory and photosensory behaviors (Bento et al., 2010; Ishita et al., 2023; Lightfoot et al., 2019; Nakayama et al., 2024; Okumura et al., 2017; Ragsdale et al., 2013; Rudel et al., 2008). Although this species has advantages for genetic manipulation as a genetic satellite model, no temporal expression tools have yet been developed.

Here, we characterized heat-inducible genes in *P. pacificus*. RNA-seq analysis revealed that thousands of genes were induced in response to heat stress. Among these genes, we found that the orthologs of *C. elegans* heat shock proteins, which are highly divergent in *P. pacificus*, were dramatically upregulated. Using the upstream regulatory region of one of these genes, we developed a heat-inducible gene expression system in this animal.

## Materials and methods

### Nematode strains and culture condition

Worm strains were cultured under standard culturing conditions (Stiernagle, 2006) at 20 °C or 15 °C in the dark (Stiernagle, 2006). Following strains were used in this paper: *P. pacificus* wildtype strain (PS312), MOK235: *Ex[PPA12242p::FLP::T2A::TurboRFP, Ppa-egl-20p::GFP]* (*excbh48*), MOK236: *Ex[PPA38778p::FLP::T2A::TurboRFP, Ppa-egl-20p::GFP]* (*excbh49*). For convenience, shorter transgene names were used for the transgenic strains as follows: *PPA12242p::TurboRFP* for MOK235 and *PPA38778p::TurboRFP* for MOK236, in following sections.

### Heat shock condition

The worms were placed on a 35-mm NGM plate seeded with *E. coli* OP50. The plates were then placed in an incubator at the temperature of interest. In the experiments not mentioned specifically, we used the following sequence of heat shock conditions: 34 °C, 1 h; 20 °C, 1 h; 34 °C, 1h. For fluorescence imaging, the worms were placed on agar pads (see *Fluorescence Imaging* section) 2 h after the last heat shock event was completed.

### RNA sequencing

Three 60 mm NGM plates with mixed-stage worms were placed in a 34 °C incubator for 2 h. After heat shock, the worms were washed with M9 buffer three times and frozen at - 80 °C. Worms without heat shock were used as control samples. Total RNA was purified using the RNeasy Mini Kit (Qiagen, 74104), and RNA sequencing libraries were prepared by BGI with the DNBSEQ Eukaryotic Strand-specific mRNA library protocol. RNA sequencing was performed by BGI using the DNBSEQ platform. Adapter sequences and low-quality reads were trimmed from the raw data. This was performed by BGI using SOAPnuke (Chen et al., 2018), followed by further trimming with Trim-Galore v0.6.6 (https://github.com/FelixKrueger/TrimGalore). The trimmed reads were then mapped to the El Paco genome assembly (Rödelsperger et al., 2017) using HISAT2 v2.2.1(Kim et al., 2019) with the --dta option. The mapped data were converted to BAM format and indexed with SAMtools v1.16 (H. Li & Durbin, 2009). Read counts were generated using StringTie v2.1.2 with the -e and -B options (Kovaka et al., 2019) based on a modified version of the El Paco V3 gene annotations, as described below. The resulting ctab files were converted to a gene count matrix in CSV format using the prepDE.py3 script (https://ccb.jhu.edu/software/stringtie/dl/prepDE.py3). Normalization and identification of differentially expressed genes (DEGs) were performed using TCC-GUI (Su et al., 2019) with R v4.4.1 using the following parameters: norm.method = “deseq2”, test.method = “edger”, iteration = 3, FDR = 0.05, and floorPDEG = 0.05.

### Restoration of Missing 3′ UTRs in a Reference Gene Annotation

The latest *P. pacificus* gene annotation, El Paco V3, incorporates extensive community-driven manual curation (Athanasouli et al., 2020; Rödelsperger et al., 2019) and contains 28,896 protein-coding genes; however, 22,741 (78.7 %) of these models lack an annotated 3′ untranslated region (3′ UTR). To restore the missing 3′ UTRs, we transferred information from three previously published annotations that shared the same El Paco genomic coordinates and included 3′ UTRs: El Paco V1 (Rödelsperger et al., 2017b), a strand-specific transcriptome assembly (Rödelsperger et al., 2016), and an Iso-Seq assembly (Werner et al., 2018). For each El Paco V3 transcript lacking a 3′ UTR, the genomic coordinates of its terminal coding sequence (CDS) in the GFF3 file served as a lookup key. This key was used to query the three source annotations for transcripts with an exactly matching terminal CDS, and their associated 3′ UTRs were retrieved. When multiple candidates were identified for a single gene, we provisionally selected the longest one. The provisional 3′ UTR was then examined for overlap with other El Paco V3 gene models on the same chromosome. If no overlap detected, the 3′ UTR was integrated into the El Paco V3 annotation without modification. If overlap was present, all candidate 3′ UTRs were pooled, and each was trimmed exon-by-exon to remove segments intersecting another gene. The longest trimmed fragment was then integrated into the El Paco V3 annotation. When trimming eliminated all candidates, we permitted the minimum necessary intersection by appending only the first exon of the candidate 3′ UTR with the least overlap. This procedure produced a consolidated set of 3′ UTR annotations while preserving the structural integrity of the existing El Paco V3 gene models.

### Phylogenetic tree of hsp-16 orthologues

The orthologs of *hsp-16* in *P. pacificus* were identified from the *P. pacificus* El Paco V3 gene annotation (Athanasouli et al., 2020), and the amino acid sequences were retrieved from Pristionchus.org (http://pristionchus.org). The amino acid sequences were aligned using MAFFT v7.525 (Katoh & Standley, 2013), and the maximum likelihood tree was generated using raxmlGUI 2.0 (Edler et al., 2021). The tree was visualized with FigTree v1.4.4 (http://tree.bio.ed.ac.uk/software/figtree/).

### Transgenic strain generation

To generate reporter lines of *PPA38778* and *PPA12242*, the genomic regions upstream of the predicted start codons of these genes (5111 bp for *PPA38778* and 1575 bp for *PPA12242*) were amplified by polymerase chain reaction (PCR) and cloned into a plasmid containing *Pristionchus*-optimized *FLP*, *T2A* sequence, and *Pristionchus*-optimized *TurboRFP*, which was linearized with FastDigest *Sma*I (ThermoFisher, FD0663), using NEBuilder (NEB, E2621). The following primers were used to amplify promoter sequences for plasmid construction: *PPA12242p*, forward: 5′-tgcctgcaggtcgacgtcccTAAATGTTCCTAATCTTGTTCTCATG-3′, reverse: 5′- actggggcatctgaaaacccTAGAGAGGGTGTACGGTAGTTC-3′; *PPA38778p*, forward: 5′- tgcctgcaggtcgacgtcccCAAATGAAATAGAGAGTCACCATATTTC-3′, reverse: 5′- actggggcatctgaaaacccTGTAGACAGCAATCGGTAGC-3′. (Uppercase, complementary bases to each locus; lowercase, homology arms for the Golden Gate reaction). The plasmids linearized with FastDigest *Hind*III (ThermoFisher, FD0504) (1 – 2 ng/µL) were injected into young adult hermaphrodites (PS312) together with genomic DNA (60 ng/µL) and injection marker plasmid pZH008 (5 ng/µL) (Han et al., 2020) digested with the same enzyme. F1 progeny were screened using a fluorescence stereoscope (Leica, M165 FC).

### Fluorescence imaging

The transgenic animals were placed on 2% agar pads. For immobilization, 2 – 5 µL of 5 mM levamisole hydrochloride (Fujifilm, 123-04641) was dropped onto the agar pads. We used levamisole because sodium azide may induce the expression of heat shock proteins in nematodes, as previously reported in *C. elegans* (Massie et al., 2003). Z-stack images of the worms were acquired using a laser scanning microscope (Zeiss, LSM 900) and Zen software. For the quantification of fluorescence intensity, the mean fluorescence intensity of the whole body was calculated for each Z slice using Fiji software (Schneider et al., 2012) for each individual, and then the mean of the fluorescence intensities in all slices for individuals was calculated. In the same experiment, the same laser intensity and gain conditions were used. The original image data are available upon request.

### Larval growth assay

For synchronization of worm developmental stages, twenty 2-day adult hermaphrodites were placed onto the plates and allowed to lay eggs for 2 h at 20 °C. After removing the adult animals, the number of eggs on each plate was counted, and the plates were placed at 15 °C. In our experiments, the worms grew into comma-stage embryos, J2, J3, and J4 larvae after 20, 75, 98, and 138 h after egg-laying, respectively. Therefore, we treated the worms with the “double heat shock,” as mentioned before at that time point. At 220 h after egg laying, the developmental stages of the worms were examined using a stereoscope (Zeiss SteREO Discovery.V20). The survival rate was calculated by dividing the number of worms alive at this time point on each plate by the number of eggs at the 0 h time point on the same plate. The number of eggs per plate ranged from 41 to 84.

#### Behavioral assay

All behavioral assays were performed at 20 °C, and the experiments were performed for at least two separate days.

### Body bending assay

Day 1 young adult hermaphrodite animals were loaded into a 96-well plate with 100 µL of M9 buffer. The number of body bends per 10 s was counted. A single body bending event was defined as a full wave-like movement of the worm’s body, such that both the head and tail returned to the same side of the body axis as they were at the initial time point. Body bending was counted three times per individual, and the average body bending was used for statistical analysis.

### Corpse assay

The corpse assay was performed as previously described (Wilecki et al., 2015). *C. elegans* victims were collected from freshly starved NGM plates and filtered through double 20 µm nylon mesh (Millipore, NY2004700). The worms were washed three times, and 2 µL of the worm pellet was placed onto empty 6-cm NGM plates. Five *P. pacificus* day 1 to day 2 adult hermaphrodites were placed on a plate with *C. elegans* larvae for 2 h. The mouth form of the predators was examined after the assay, and the number of corpses per Eurystomatous animal was calculated by dividing the number of corpses per plate by the number of Eu animals in the plate.

#### Mouth form ratio

The mouth form ratio was examined as previously described (Ragsdale et al., 2013). Briefly, animals on day 2 to 3 were anesthetized with 0.3% sodium azide, and the mouth form was observed at 400x magnification using a DIC microscope.

#### Statistical analysis

R software and Microsoft Excel were used for statistical analysis. The type of statistical test, significance symbols, and n number are represented in the figure legends.

#### Data Availability

All materials generated in this study, including plasmids and worm strains, and the data underlying this article are available upon request from the corresponding author. RNA-sequencing data are available in DDBJ at https://www.ddbj.nig.ac.jp/index-e.html, and can be accessed with PRJDB 35585.

## Results

### Heat induced changes of expression level in thousands of genes

To identify the genes highly induced by heat shock in *P. pacificus*, we performed RNA sequencing analysis to compare the global expression levels of genes between normal and heat-shock conditions. Because heat induction of gene expression was successful at 34 °C for 1 h or 33 °C for 2 h in *C. elegans* (Davis et al., 2008; Hubbard, 2014; Voutev & Hubbard, 2008), we used heat shock conditions with a higher temperature and longer time (that is 34°C for 2 h) in our RNA-seq study. To improve the accuracy of read count quantification using the current version of gene annotation (El Paco V3 annotation) (Athanasouli et al., 2020), whose 3′ untranslated regions (3′ UTRs) were not annotated in many genes, we generated a modified annotation file by extending 3′ UTRs using three previously published annotation files (see Materials and Methods section). In the comparison of gene expression between control samples and heat-shocked animals, we found that as many as 7815 genes were differentially expressed between the groups under a 5% FDR condition (Figure 1A).

**Figure 1.**
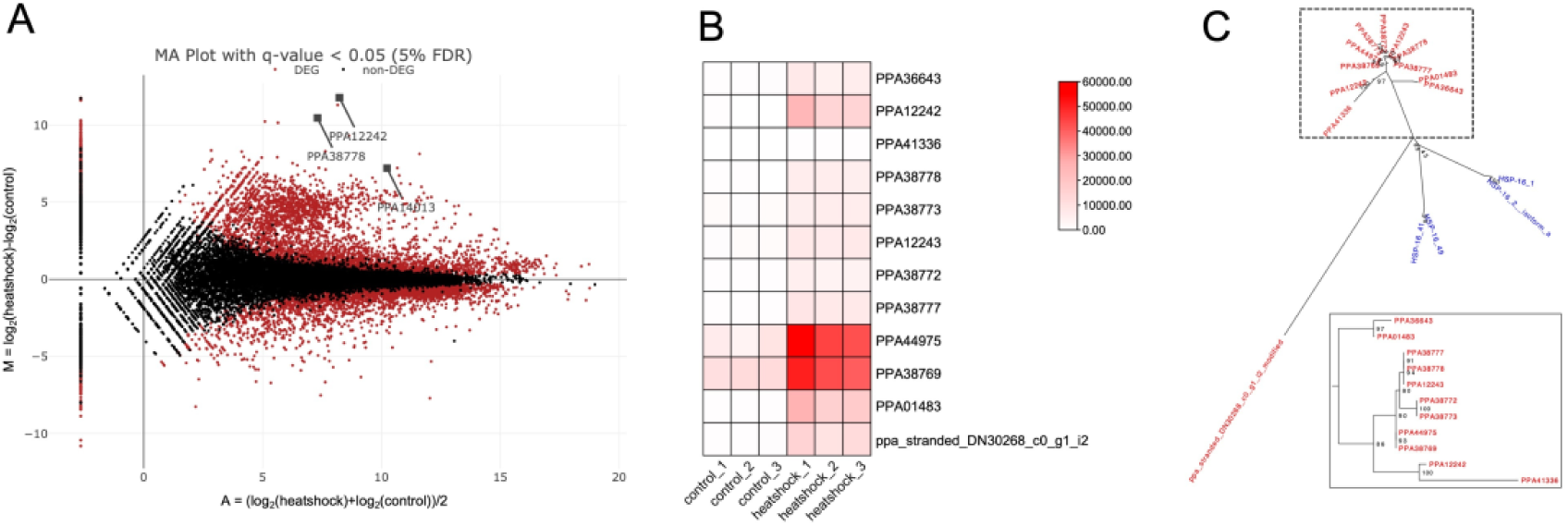
Twelve orthologs of *hsp-16* were induced upon heat shock. (A) MA plot of the RNA-seq data of global gene expression between the control and heat-shocked animals. Genes that were significantly upregulated or downregulated are highlighted in red (FDR-adjusted P < 0.05). HSP orthologs ranked within the top ten DEGs are highlighted with gray squares. (B) Heatmap representing the normalized read count of 12 orthologs of *hsp-16* found in the *P. pacificus* genome. (C) Unrooted phylogenetic tree of *hsp-16* paralogs found in *P. pacificus* and *C. elegans*. Genes in *P. pacificus* and *C. elegans* are shown in red and blue, respectively. The left-bottom tree is a magnified tree of genes enclosed in a dashed-lined rectangle.

### Orthologs of hsp-16 were upregulated in response to heat

The top ten differentially expressed genes (DEGs) included three genes orthologous to small heat shock proteins in *C. elegans* (Table 1). Two of them (*PPA38778* and *PPA12242*) encoded the orthologs of *hsp-16.41*, whose promoter is used for ubiquitous gene induction by heat shock in *C. elegans* (Voutev & Hubbard, 2008). In the current version of gene annotation (Athanasouli et al., 2020), we found 12 paralogs of *hsp-16* in the *P. pacificus* genome, all of which were robustly upregulated upon heat shock (Figure 1B). To determine whether there are one-to-one orthologs of *hsp-16* genes in *C. elegans*, we constructed a maximum likelihood tree using the protein sequences of *hsp-16* orthologs in *P. pacificus* and *C. elegans* (Figure 1C). In *C. elegans* genome, there are six *hsp-16* paralogs (*hsp-16.1*, *hsp-16.2*, *hsp-16.11*, *hsp-16.41*, *hsp-16.48*, and *hsp-16.49*) encoding four proteins (HSP-16.1, HSP-16.2, HSP-16.41, HSP-16.49); *hsp-16.11* and *hsp-16.48* encode same polypeptides as *hsp-16.1* and *hsp-16.49*, respectively. The *hsp-16* genes were clustered within the species, and there appeared to be no one-to-one orthologs of *hsp-16* genes between *C. elegans* and *P. pacificus*. All but one of the 12 paralogs in *P. pacificus* represented very short branch lengths, suggesting that these paralogs arose rapidly during evolution.

**Table 1.** Top ten DEGs upregulated upon heat shock.

| Rank | Gene name | *C. elegans* ortholog | GO term |
| --- | --- | --- | --- |
| 1 | PPA02457 | dpy-5_best | Collagen,<br>Col_cuticle_N |
| 2 | PPA38778 | hsp-16.41_best | HSP20,<br>ArsA_HSP20 |
| 3 | ppa_stranded_DN24775_c0_g2_i1 | cdr-6_BRH | GST_C_2,<br>GST_C_6,<br>GST_N_4 |
| 4 | ppa_stranded_DN19351_c0_g2_i1 | cdr-2_best | GST_C,<br>GST_C_2,<br>GST_C_6,<br>GST_N_4 |
| 5 | PPA12242 | hsp-16.41_best | HSP20 |
| 6 | ppa_stranded_DN16198_c0_g1_i1 | cyp-33C5_BRH | p450 |
| 7 | PPA42171 | col-3_best | Collagen,<br>Col_cuticle_N |
| 8 | PPA40518 | col-3_best | Collagen,<br>Col_cuticle_N |
| 9 | PPA14013 | hsp-25_best | HSP20 |
| 10 | PPA33590 | col-3 | Collagen,<br>Col_cuticle_N |

### Heat induced PPA12242 gene expression in several tissues

As RNA-seq data revealed that *PPA38778* and *PPA12242* were robustly expressed under heat shock conditions, while they were almost not expressed at normal culture temperature, we generated transgenic lines expressing TurboRFP reporter under the promoter of *PPA38778* or *PPA12242*. We examined the expression of RFP in animals carrying the transgene with or without heat shock.

In the *PPA38778p::TurboRFP* strain, RFP was not expressed in any tissues under standard culture conditions (20 °C) or heat-shock conditions (34 °C, 2 h) (Figure S1A, B), suggesting that the upstream sequences of *PPA38778* are not sufficient to induce gene expression in response to heat shock events. In the *PPA12242p::TurboRFP* strain, however, normal culturing temperature (20 °C) resulted in the expression of the RFP reporter in some of the head and tail neurons, as well as non-neuronal cells in the head (Figure S2A). To reduce the leaky expression of transgenes, we cultured the transgenic animals at 15 °C, which is cooler than the standard culturing condition (Stiernagle, 2006). This relieved the unwanted expression of RFP, and only a pair of head neurons expressed RFP (Figure S2B). Thus, we used this growth temperature for subsequent analyses.

Unexpectedly, the RFP signal was not visible in *PPA12242p::TurboRFP* animals after 34 °C, 2 h of heat shock, which was used for the RNA-seq analysis. Therefore, we attempted to determine the optimal heat-shock conditions for the efficient induction of genes with the *PPA12242* promoter. We examined higher temperatures, multiple heat shock events, pre-heat and heat shock events, and sodium azide treatment, which is known to induce heat shock proteins in *C. elegans* without heat shock (Massie et al., 2003) (Figure S2C). Among the conditions we tried, the “double heat shock” condition with two heat shock events at 34 °C for 1 h each franking 1 h at 20 °C cooling down worked well (Figure 2A); the RFP fluorescence level was three times higher than the basal level of fluorescence, which was derived from autofluorescence in the intestine (Figure S2C).

**Figure 2.**
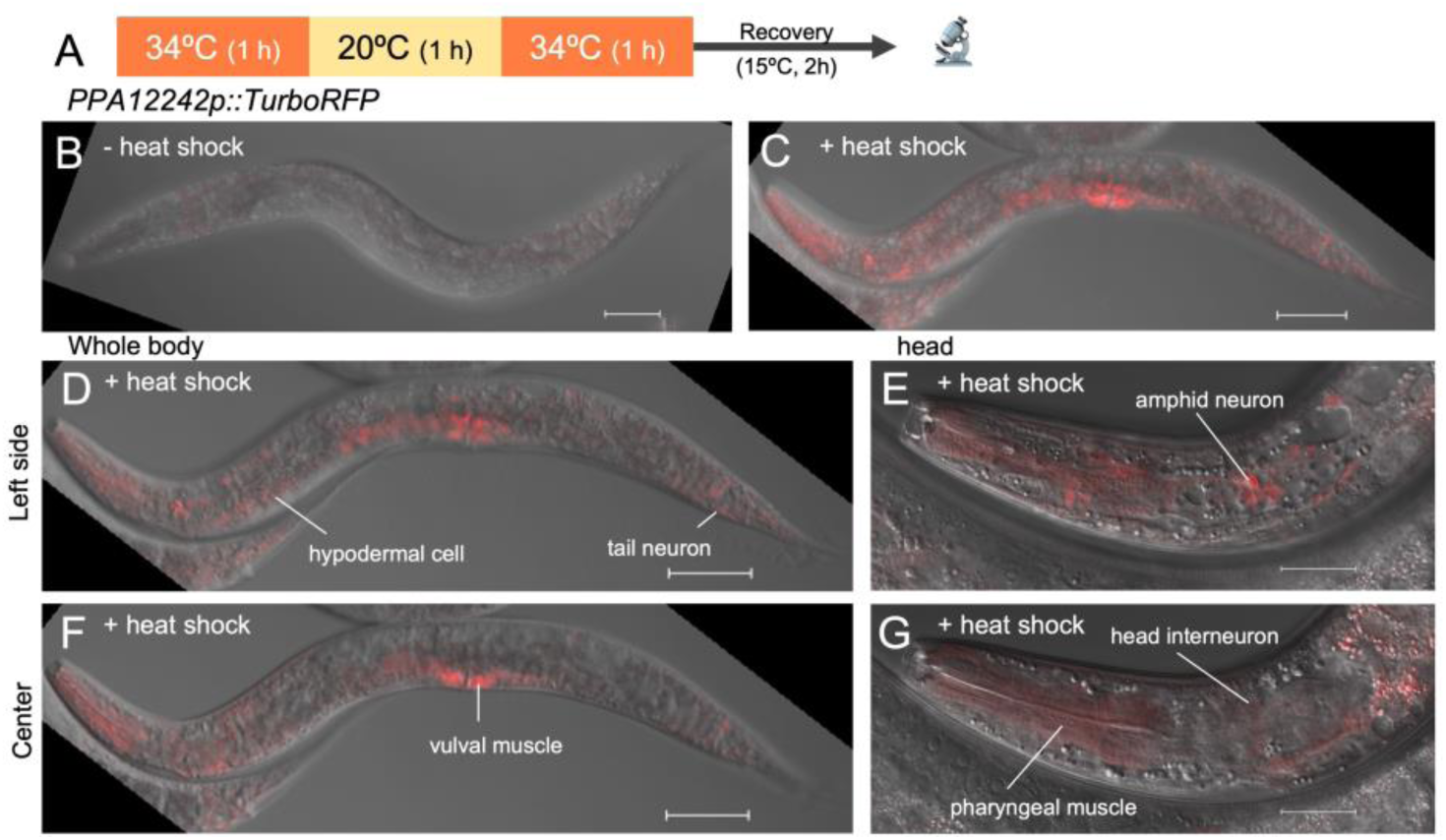
Expression patterns of RFP under the *PPA12242* promoter following heat shock. (A) Schematic representation of the heat shock treatment schedule. (B, C) Maximum projection images of *PPA12242p::TurboRFP* on adult day 1. RFP fluorescence was not obvious in the worms cultured at 15°C (B), but it was observed in the heat-shocked animals (C). Scale bars, 50 µm. (D – G) Single-plane images of *PPA12242p::TurboRFP* animals. Planes focusing on the left side (D, E) and center (F, G) of the worms are shown. Magnifications of 200 × and 400 × were used for (D, F) and (E, G), respectively. Scale bars, 50 µm and 20 µm for (D, F) and (E, G), respectively. Note that this animal is the same as that in (C).

Heat shock events induced RFP expression in several tissues in *PPA12242p::TurboRFP* young adult hermaphrodites; RFP fluorescence was observed in vulval tissues, hypodermis, head neurons, including amphid neurons, tail neurons, and pharyngeal muscles (Figures 2B – G). With increased brightness and contrast, the RFP signal was observed in most tissues, excluding the germline cells.

Taken together, these results indicate that the *PPA12242* promoter can be used as a heat-shock promoter in *P. pacificus*.

### PPA12242 was expressed intensely in larval stages

Next, to validate the use of *PPA12242p* at various developmental stages, we examined the expression of RFP in *PPA12242p::TurboRFP* animals in three larval stages, together with the young adult stage (Figures 3A, B). In comparison to the mean fluorescence intensity among the animals, we found that the mean RFP expression level was highest in the J3 larval stage and lowest in the adult stage (Figure 3B). In J3 animals, the RFP signal was obvious in most tissues, excluding germline precursor cells, which are generally inactive in transgene expression in this species (Figure 3C). In J4 animals, RFP expression was intense in developing tissues, including the vulva, hypodermal tissue, arcade cells, and pharyngeal muscles (Figure 3A). Although there seem to be substantial differences in the expression level of RFP, *PPA12242p* is applicable at all developmental stages.

**Figure 3.**
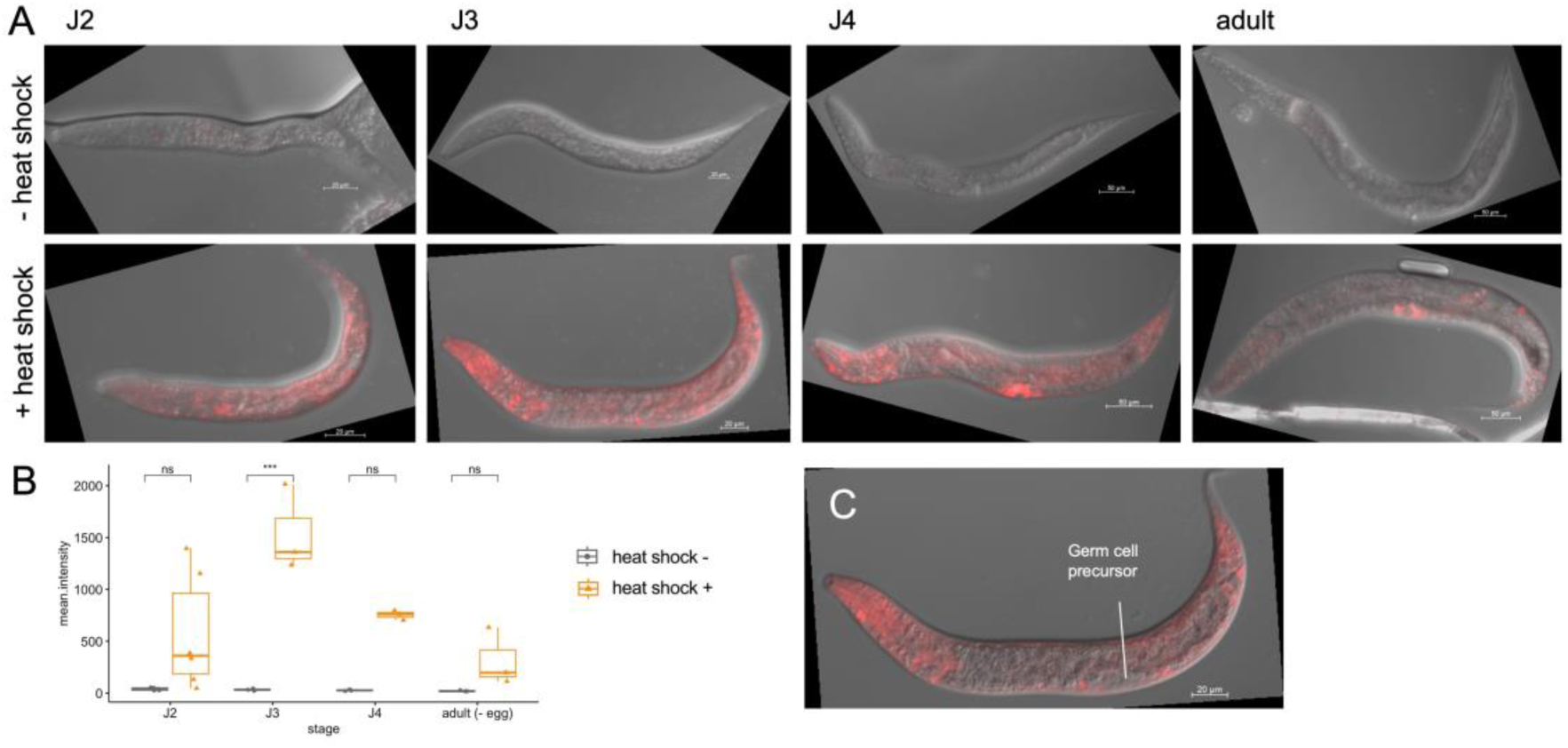
Expression patterns of RFP under the *PPA12242* promoter following heat shock during each developmental stage. (A) Maximum-projection images of *PPA12242p::TurboRFP* animals at different developmental stages before (upper panels) and after (lower panels) heat shock treatment. Scale bars, 20 µm for J2 and J3 animals; 50 µm for J4 and adult animals. (B) Quantification of fluorescence intensity in *PPA12242p::TurboRFP* animals at different developmental stages before and after heat-shock treatment. Two-way ANOVA with Tukey’s multiple comparison test. ns, P ≥ 0.05. *** P < 0.001. (C) Single-plane image of *PPA12242p::TurboRFP* J3 animal after heat shock treatment, focusing on the center plane. Scale bar, 20 µm.

### Specific larval stage was vulnerable to heat stress

Heat shock events are stressful for organisms and alter their physiological state. To determine whether heat shock events alter the development and behavior of *P. pacificus*, we examined several phenotypes after heat shock events.

First, we examined the survival rate and developmental delay in heat-shocked animals. Due to a development delay of up to a week when *P. pacificus* is cultured at 15 °C, we scheduled the heat-shock event at 20, 75, 98, and 138 hours after egg laying, corresponding to the embryo, J2, J3, and J4 stages, respectively. (Figure 4A). Among the animals that did not undergo heat shock events, 88.9% survived 196 hours post-egg laying. The survival rate of heat-shocked animals at the J2 and J4 stages was not significantly altered compared to that of animals without heat shock. In contrast, heat shock events at the embryo and J3 stages significantly reduced the survival rate. Especially, the survival rate was dropped to 0.74% when the worms were heat-shocked at the embryonic stage. This prevented us from characterizing the phenotypes in subsequent analyses (Figure 4B). This result is different from that of a previous *C. elegans* study which showed that the embryo of this species is highly tolerant to heat shock events owing to the protective effect of an embryonic heat shock protein (Fleckenstein et al., 2015).

**Figure 4.**
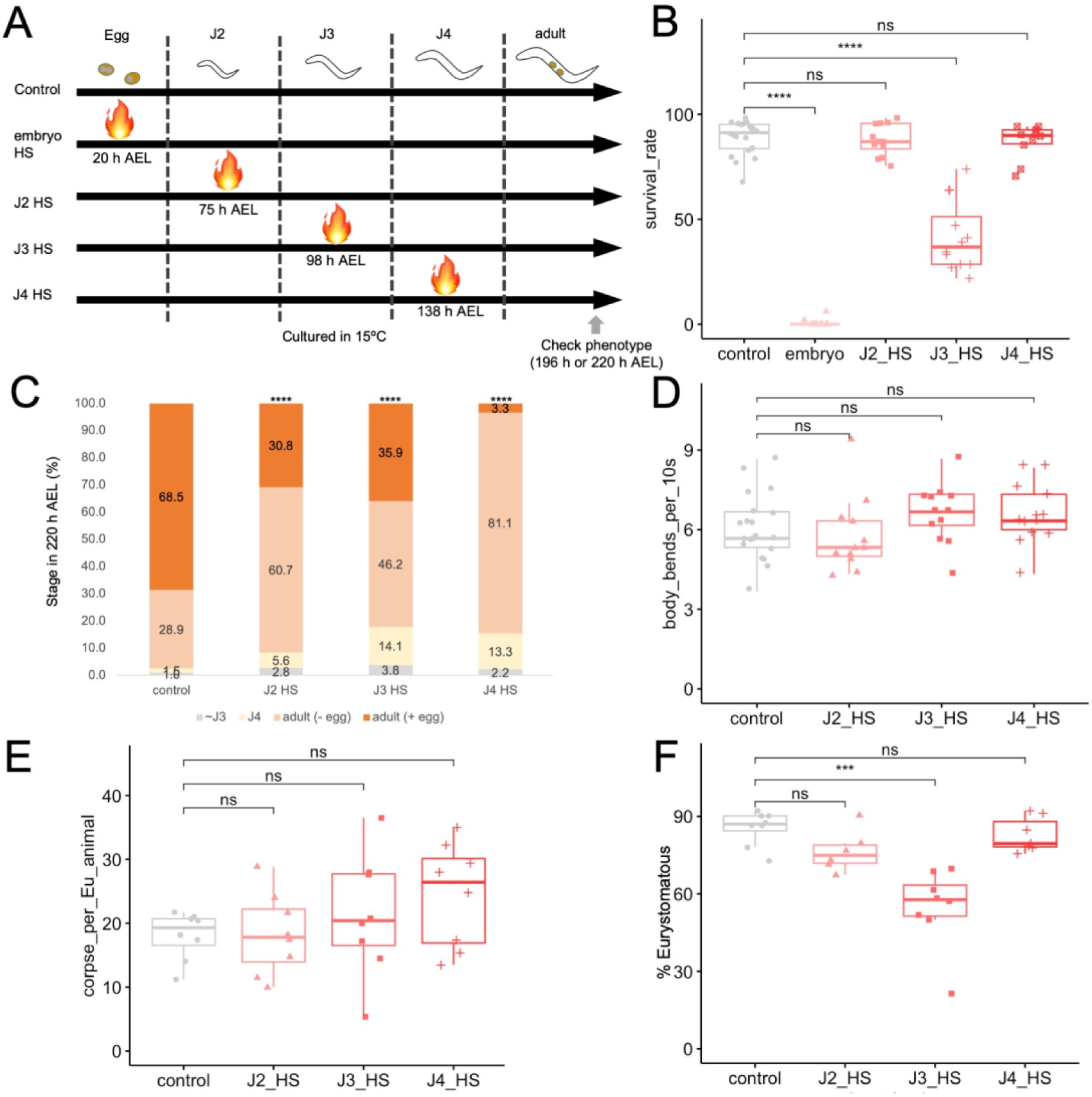
Phenotypic characterization of worms that experienced heat shock events during larval stages. (A) Schematics of the heat shock schedules and phenotyping. HS, heat shock. AEL, after egg laying. (B) Survival rates of wild-type animals subjected to heat shock at each developmental stage. n=20, 11, 12, 12, and 10 plates for control (without heat shock), embryo, J2, J3, and J4 with heat shock, respectively. Kruskal-Wallis test with Steel’s multiple comparison test. ns, P ≥ 0.05. **** P < 0.0001. (C) Proportions of developmental stages at 220 h after egg laying. Numbers in each column represent the proportion of each developmental stage (%). The proportions of each stage are shown in gray, light yellow, light orange, and orange for younger than J3 (∼ J3), J4, adults without eggs (adult (- eggs)), and adults with eggs (adult (+ eggs)), respectively. n = 197, 107, 78, 90 for control (without heat shock), J2 HS, J3 HS, and J4 HS, respectively. Chi-square tests were used to compare each column with the control. **** P < 0.0001. (D) Number of body bendings per 10 s in M9 buffer. n = 21, 12, 12, 13 for control, J2 HS, J3 HS, J4 HS, respectively. One-way ANOVA. ns, P ≥ 0.05. (E) Number of corpses per eurystomatous animal after 2 h of corpse assay. n = 8 for all heat shock conditions. One-way ANOVA. ns, P ≥ 0.05. (F) Proportion of eurystomatous animals. n = 8, 6, 8, 7 for control, J2 HS, J3 HS, J4 HS, respectively. One-way ANOVA with Dunnett’s multiple comparison test. ns, P ≥ 0.05. *** P < 0.001.

We also examined whether heat shock delayed developmental speed. At 220 h after egg-laying, 97.5% of the control animals without any heat shock events grew into the adult stage, and 68.5% of them were reproductive adults. However, heat shock events during the larval stages delayed development. For example, the proportions of individuals reaching the adult stage were 91.5%, 82.1%, and 84.4% for those that experienced heat shocks during the J2, J3, and J4 stages, respectively (Figure 4C).

Next, we performed a body-bending assay under liquid conditions to observe the locomotor activity of heat-shocked animals. In *C. elegans*, locomotory behavior is commonly used to assess nervous system defects (Dimitriadi & Hart, 2010; Hornsten et al., 2007; J. Li et al., 2016; Zhang & Chen, 2023). The average number of body bendings per 10 s was 6 in the control animals, which was consistent with a previous report (Ishita et al., 2021). Heat-shocked animals did not significantly alter the number of body bends (Figure 4D).

As mouth-form dimorphism and predatory feeding behavior are unique and well-characterized phenotypes of *P. pacificus*, we examined these traits in animals subjected to heat shock during the larval stages. To measure predatory feeding events, we used a corpse assay (Lightfoot et al., 2016; Wilecki et al., 2015) (See Materials and Methods). In wild-type animals, predatory killing within 2 h per Eurystomatous animal was 18.1 on average (Figure 4E). The number of corpses per Eurystomatous animals that were heat-shocked during the larval stages did not significantly differ from that of the control animals (Figure 4E), suggesting that predatory feeding behavior is robust to heat shock events during the larval stages. However, in the case of mouth-form ratio, the specific larval stage was sensitive to heat shock events. The control PS312 animals exhibited more than 80% Eurystomatous animals, consistent with previous studies (Ragsdale et al., 2013). Compared with the control animals, worms heat-shocked at the J3 stage showed a decreased proportion of Eurystomatous animals to 54.5 % on average (Figure 4F). The mouth-form ratio in animals heat-shocked at the J2 and J4 stages was not significantly altered.

Taken together, these data suggest that some developmental traits are vulnerable to heat-shock events, especially during the J3 stage, but other traits, including predatory feeding behavior and locomotory behavior, are robust to developmental heat stress.

## Discussion

Heat stress is one of the most ubiquitous stresses in all organisms. In this study, we identified genes that were upregulated in response to heat stress. Specifically, we found that some heat shock protein orthologs are highly expressed upon heat shock by hundreds or thousands of folds, suggesting their functional conservation, even after diversification. With a typical heat shock promoter structure, *PPA12242* was induced by an artificial heat shock event, indicating the utility of this promoter as a heat-inducible gene expression system. Phenotypic characterization of heat-shocked animals revealed that some developmental traits are vulnerable to heat stress at specific developmental stages, while most of the traits we tested did not show significant differences after short-term heat shock events. This study establishes a temporal gene expression system induced by heat shock using the promoter of a heat shock protein, offering the first temporal expression system in *P. pacificus*.

Heat shock proteins are a highly conserved gene family among organisms; however, some of them are duplicated in certain species (Gong & Golic, 2004; Hu et al., 2022; Nikolaidis & Nei, 2003; Obuchowski et al., 2019). In this study, we found that the *hsp-16.41* paralogs were highly divergent in *P. pacificus*. RNA-seq analysis indicated that all 12 paralogs were induced by heat treatment, suggesting the conservation of the heat-protective function of these molecules. Transcriptional reporter analysis of the *PPA12242* promoter suggests that there are differences in the expression level of this gene among developmental stages and cell types, while heat shock response itself is required for all stages and cell types. Other diversified orthologs may compensate for the expression of *PPA12242* in other stages and cell types, as explained for *C. elegans hsp-16.2* and *hsp-16.41* paralogs (Stringham et al., 1992).

Curiously, not all *hsp-16.41* paralogs had typical heat-shock factor motifs with a TATA box upstream of their coding sequences, although they were robustly induced by heat treatment. The *PPA38778* reporter showed no activation after heat shock, suggesting that its expression may be regulated by elements other than the upstream promoter. In mammalian cells, most heat shock response genes are induced independently of heat shock factors, and many are acutely induced by heat shock (Mahat et al., 2016). Although the exact transcriptional mechanism of *PPA38778* was not clarified in this study, the heat-induction mechanism characterized in other genes, such as the SRF transcription factor (Mahat et al., 2016), in combination with more distal elements or genome-wide epigenetic changes, might be responsible for the induction of this gene.

The difference in the expression level of *PPA12242* among developmental stages may be associated with vulnerability to heat stress at specific developmental stages. The expression level of RFP driven by the *PPA12242* promoter was highest during heat shock in the J3 stage. Heat stress at this stage increased lethality compared to other larval and young adult stages, suggesting that the J3 stage might be vulnerable to heat stress and that heat shock proteins might be highly induced to protect cells from the deleterious effects of heat. This is comparable to a previous study in *C. elegans*, which showed that the knockdown of heat shock factor induces developmental arrest in the L2/L3 stage when worms are cultured at high temperatures (Walker et al., 2003). It would be interesting to compare the transcriptome changes in J3 heat shock conditions and those in other heat shock conditions to investigate the cause of vulnerability at this stage.

In *P. pacificus*, a heat shock event at the J3 stage also altered the mouth form ratio in the adult stage. This result is consistent with a previous report indicating that the critical period of mouth-form determination is around the J3-J4 molt in this species (Werner et al., 2023). The mouth form ratio of this species is affected by environmental conditions, including starvation, temperature changes, and culture conditions, which would induce heat shock proteins (Bento et al., 2010; Lenuzzi et al., 2021; Werner et al., 2017). There may be some direct or indirect association between heat shock proteins and mouth-form determination, which will be elucidated in future studies.

Taken together, this study demonstrates heat-inducible gene expression using an endogenous heat shock promoter in *P. pacificus*. This would offer one of the first examples of a conditional gene expression system in this species; however, there are some limitations to this system. First, leaky expression of genes with the heat shock promoter is inevitable in some cells, even at lower temperatures than the standard culture conditions. Similar problems are often observed in other model organisms, but this can be improved by using additional gene regulatory elements, such as polycomb response elements (Akmammedov et al., 2017). Second, because heat stress disturbs many biological processes and affects the phenotypes of interest, including mouth-form determination, alternative gene or protein expression strategies, such as the Q system, Tet-On/Off system, and GeneSwitch, should also be developed for this species (Bello et al., 1998; Osterwalder et al., 2001; Potter et al., 2010). Third, the heat shock promoter used in this study could not induce gene expression in some cell types. The expression in some cell types could be improved by using other heat shock promoters in combination with *PPA12242p.* However, in germline cells, other technologies for transgenesis, including gene bombardment (Namai & Sugimoto, 2018) or other regulatory strategies, such as PATC repeats found in *C. elegan*s (Aljohani et al., 2020), would be required. With these new genetic tools, the developmental stage-specific function of genes can be unraveled in this species, accelerating the comparative genetic study of model nematode systems.

## Acknowledgements

We would like to thank Dr. Ziduan Han and Dr. Ralf J Sommer (Max Planck Institute for Developmental Biology) for providing the optimized fluorescent plasmids. We thank all the members of the Chihara Laboratory and Dr. Kohta Yoshida (Niigata University) for their helpful support in this study. This work was supported by the JST FOREST Program (Grant Number JPMJFR214M), Research grants in the Natural Sciences (The Mitsubishi Foundation), Narishige Zoological Science Award, Yamada Science Foundation, and JSPS KAKENHI (Grant Number JP25K02318) to M.O.; JST SPRING (Grant Number JPMJSP2132) to H.H.; JSPS Research Fellows (Grant Number 22J13816) to Y.I.

## Author contribution

Conceptualization: Y.I. and M.O.; methodology: Y.I. and M.O.; investigation: Y.I., H. H. and M.O.; writing—original draft: Y.I.; writing—review and editing: H.H., T.C., and M.O.; supervision: T.C. and M.O.; funding acquis sition: Y.I., H.H., and M.O. All authors have approved the final version of the manuscript.

