## Supplemental Figures for "Characterization of heat shock protein expression and its application to temporal gene expression in the nematode *Pristionchus pacificus*"

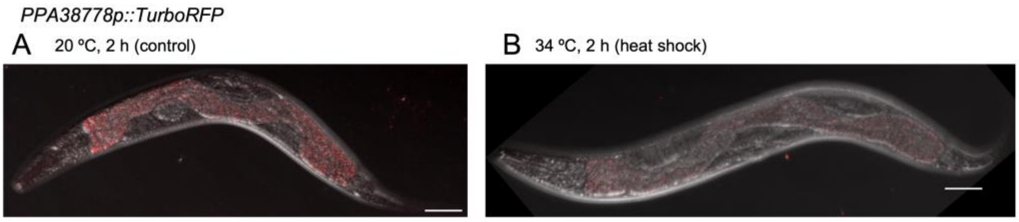


**Figure S1. RFP expression in *PPA38778p::TurboRFP*.**

(A, B) RFP expression before (A) and after (B) heat shock treatment in *PPA38778p::TurboRFP*. Scale bars, 50 μm. Red fluorescence in the intestine is autofluorescence.

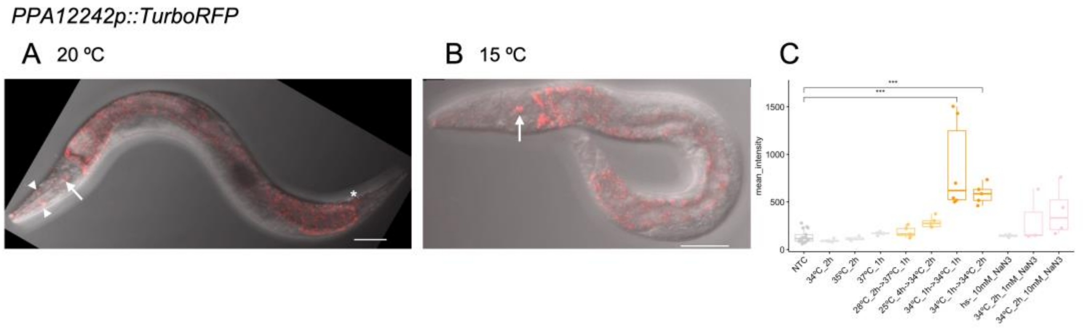


**Figure S2. Optimization of culture and heat-shock conditions.**

(A) RFP expression in head neurons (arrow) and other cells including head hypodermal cells (arrowhead) and tail neurons (asterisk) in *PPA12242p::TurboRFP* cultured at 20 °C. Scale bar, 50 µm. Red fluorescence in the intestine is autofluorescence.

(B) RFP expression in a pair of head neurons (arrow) in *PPA12242p::TurboRFP* cultured at 15 ºC. Scale bar, 50 µm. Red fluorescence in the intestine is autofluorescence.

(C) Quantification of fluorescence intensity levels in *PPA12242p::TurboRFP* animals under various heat-shock conditions. One-way ANOVA with Dunnett's multiple comparison tests. *** $P < 0.001$. Box plots without significance symbols are $P \geq 0.05$.
